# RegDesign-Bench: Benchmarking computational methods for cell-type-specific regulatory DNA design

**DOI:** 10.64898/2026.10.18.759890

**Authors:** Jiaying Xu, Zekun Guo, Cuiting Zhong, Qijin Yin, Yuan He, Zeyu Chen, Hongyi Xin, Peiran Jin, Pan Deng

**Affiliations:** Global College, Shanghai Jiao Tong University, Dongchuan Road 800, 200240, Shanghai, China; Zhongguancun Academy, 100094, Beijing, China; Zhongguancun Institute of Artificial Intelligence, 100094, Beijing, China; Department of Cell Biology, MOE Engineering Research Center of Regenerative Medicine, School of Basic Medical Sciences, Peking University Health Science Center, Beijing,China; School of Future Technology, Shanghai Jiao Tong University, Dongchuan Road 800, 200240, Shanghai, China

## Abstract

The de novo design of regulatory DNA sequences that drive cell-type-specific gene expression could enable precise gene control in synthetic biology, cell engineering and gene therapy. Various computational methods have been developed for this task, yet systematic comparisons that guide method selection for different design objectives remain limited. Here, we introduce RegDesign-Bench, a standardized framework for evaluating cell-type-specific cis-regulatory element design through a comprehensive set of computational analyses. Among the methods tested, strong predicted activity was not consistently accompanied by cell-type-selective accessibility or prominent cell-type-associated motif signals. Similarity to reference sequence distributions also did not ensure high predicted target-cell activity or specificity. These findings demonstrate that favourable performance on individual measurements does not necessarily extend across regulatory properties and library characteristics, revealing application-dependent strengths and limitations of current design methods. Overall, RegDesign-Bench offers a general-purpose computational framework for comparing and selecting methods to design regulatory DNA that drives cell-type-specific gene expression.

## 1 Introduction

Cis-regulatory elements (CREs) control where, when and how strongly genes are expressed[1–3]. Their activity depends on both the underlying DNA sequence and the cellular context, allowing the same element to drive different levels of expression across cell types[2, 4, 5]. Engineering these elements offers a way to direct gene expression to selected cell populations, with applications in functional studies[6–8], cellular engineering[9–11] and therapeutic gene delivery[12–14]. For these applications, the central design challenge is to achieve high activity in the intended cell type while maintaining low activity in relevant non-target cells.

Computational approaches offer a promising route to this goal. Recent studies have designed cell-type-, cell-state- and tissue-specific regulatory elements across diverse cellular and organismal settings by exploiting differences in regulatory activity or chromatin accessibility between target and non-target populations[15–20]. Motif-based approaches construct regulatory sequences from interpretable combinations of transcription factor binding sites, sometimes incorporating models of motif interactions or model-guided motif placement[15, 21, 22]. Data-driven approaches use predictive models to guide sequence optimization[16–20] or train generative models to produce sequences with desired cell-type-specific regulatory properties[15, 23]. Individual frameworks can combine these strategies by integrating interpretable motif rules with learned sequence-to-function models. Although these methods pursue related design objectives, their original studies differ in training data, biological systems, optimization objectives and experimental validation schemes, complicating direct comparisons across studies. Existing head-to-head comparisons provide evidence within particular evaluation settings[17, 23], but do not by themselves establish a general ranking of methods across biological contexts and application requirements. Systematic evaluation under matched design tasks and validation conditions is therefore needed to clarify their relative strengths and limitations and guide method selection.

To this end, we developed a multidimensional framework, RegDesign-Bench, that evaluates diverse and complementary properties of regulatory sequences designed by different methods. First, functionality assesses activity in the intended cell type and specificity relative to background cell types. Second, biological plausibility examines chromatin-accessibility signals[24] and cell-type-relevant regulatory motifs[25, 26], providing complementary computational evidence of regulatory potential. Third, sequence diversity quantifies variation among generated sequences and their similarity to reference sequences. Finally, computational efficiency measures the time and memory required for sequence generation. Together, these properties provide a more comprehensive view of each method’s performance and reveal differences that would be obscured by any single metric.

We applied RegDesign-Bench to eight representative data-driven methods[17, 23, 27–32] generating sequences for human cell types. The comparison revealed distinct strengths across methods, with no single approach dominating all evaluated dimensions. For example, FastSeqProp [29] and DNA-Diffusion [23] performed strongly on functionality- and biological-plausibility-related measurements, while D3 (Convolutional) [31] and TACO [32] achieved the highest computational-efficiency ranks under standardized hardware conditions. By establishing where these methods excel and where their limitations lie, our benchmark supports method selection according to application-specific design objectives and provides a reproducible basis for evaluating future regulatory-sequence generators.

## 2 Results

### 2.1 RegDesign-Bench overview

RegDesign-Bench provides a common evaluation workflow for comparing methods that design cell-type-specific cis-regulatory elements (CREs) (Fig. 1a). The design objective is to generate sequences with high regulatory activity in a specified cell type (the target cell type) and low activity in the other evaluated cell types (the background cell types). For each method, the benchmark evaluates the sequence library generated for each target cell type using a common set of evaluation criteria, enabling direct comparison of method performance.

**Fig. 1:**
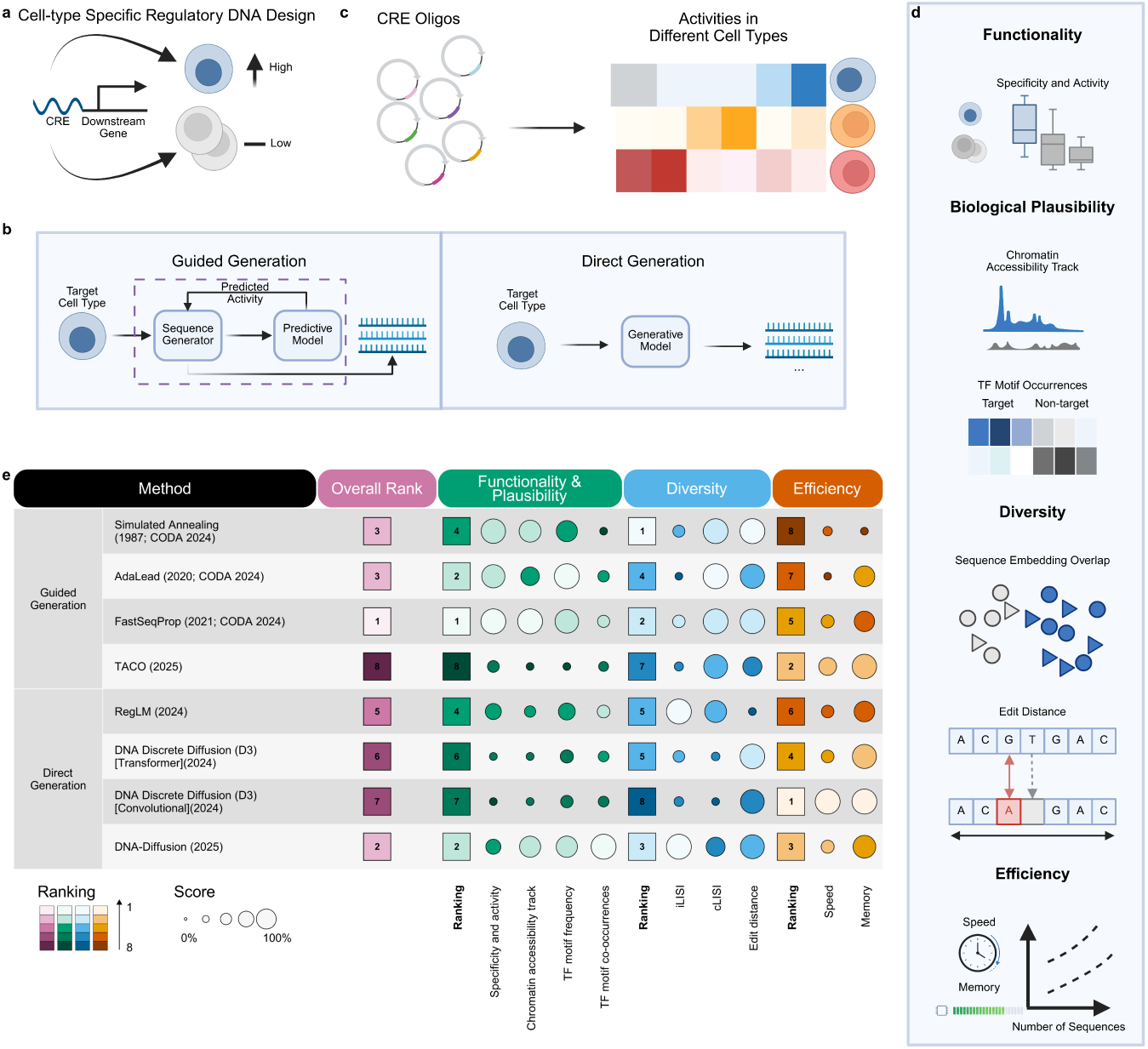
Computational strategies and benchmark for cell-type-specific regulatory DNA design. **a,** CREs are designed for high target-cell activity and low non-target-cell activity. **b,** Guided generation uses predictive or reward feedback during search or generator fine-tuning. Direct generation samples from a target-conditioned model. **c,** Published experimental MPRA data provide sequence–activity references. Heatmap rows represent cell types and columns represent CRE sequences [17]. **d,** Metrics cover functionality and biological plausibility, sequence diversity and computational efficiency. **e,** Eight methods are compared. Circle size represents the min–max-normalized metric score, colour represents metric rank, and squares show dimension and overall ranks. Higher normalized scores indicate better performance after accounting for metric direction. Rank sums determine aggregate ranks. DNA-Diffusion lacks SK-N-SH generation: activity, DNase signal area gap, both motif metrics and edit-distance rankings use metric-specific SK-N-SH mean imputation from the other seven generators; LISI uses multiplicative reference-factor adjustment (Methods 5.4.2). Created in https://BioRender.com.

We applied this workflow to eight methods spanning guided and direct sequence generation (Fig. 1b).

Guided-generation methods use predicted regulatory activity or reward signals to guide sequence design. Simulated annealing, AdaLead [28] and FastSeqProp [29] iteratively optimize candidate sequences through stochastic search, adaptive evolutionary search and gradient-based optimization, respectively. For these three methods, we retained Malinois [17], the activity predictor supplied with the original CRE-design implementation, to guide sequence optimization. TACO [32] instead uses reinforcement learning to fine-tune a pretrained RegLM [30] generator. We retained TACO’s original reward-model setup, which combines predicted regulatory activity with transcription-factor-binding-site rewards to guide model fine-tuning. These predictors and rewards guide sequence design; a separate predictor is applied during evaluation.

Direct-generation methods learn sequence distributions conditioned on the desired cell type or activity profile and sample candidate sequences from these distributions. RegLM [30] uses autoregressive generation, producing each nucleotide based on the specified activity labels and the preceding nucleotides. DNA-Diffusion [23] and DNA Discrete Diffusion (D3) [31] use diffusion models, progressively transforming noisy inputs into regulatory sequences through a learned denoising process. We evaluated two D3 variants, D3 Transformer and D3 Convolutional, which use different neural network architectures for denoising and were treated as separate methods in the benchmark.

We used publicly released checkpoints and followed the original method-specific settings, with benchmark-specific adaptations described in Methods 5.2. Therefore, generation was restricted to the cell-type conditions supported by these implementations. Seven methods used generators or optimization predictors trained on the published massively parallel reporter assay (MPRA) dataset spanning K562, HepG2 and SK-N-SH[17]. DNA-Diffusion instead was trained on sequences from DNase I-hypersensitive sites (DHSs) in K562, HepG2, GM12878 and hESCT0[23, 24]. To cover the generation conditions supported by most methods, we focused the benchmark on K562, HepG2 and SK-N-SH. Each cell type served in turn as the target cell type, with the remaining two serving as background cell types. For each method, we generated 1,000 sequences for every supported target cell type.

The generated libraries then underwent a shared downstream evaluation covering functionality, biological plausibility, sequence diversity, and computational efficiency. We used the same activity predictor, regulatory-feature analyses and sequence-comparison procedures across methods, and measured generation throughput and peak memory consumption under a common hardware protocol. Individual metric results were retained alongside dimension-specific and overall rankings, calculated by summing the corresponding metric ranks (Methods 5.4.1). For DNA-Diffusion, metrics affected by the absence of an SK-N-SH-targeted library were adjusted individually (Methods 5.4.2), while efficiency measurements were left unchanged.

The comparison revealed distinct strengths across methods, with no single approach leading every evaluation dimension. FastSeqProp ranked first overall, placing first in functionality and biological plausibility and second in sequence diversity. DNA-Diffusion ranked second overall under the coverage-adjusted aggregation, with corresponding ranks of second and third in these two dimensions. However, the coverage-adjusted values provide only an approximate basis for comparison, and its aggregate ranking should not be interpreted as directly evaluated performance across all three cell types. Simulated annealing achieved the highest diversity rank, although it had the lowest computational-efficiency rank. D3 Convolutional and TACO led computational efficiency, ranking first and second, respectively. These overall profiles establish the principal differences among methods, which we examine through the individual evaluations below.

### 2.2 Predicted cell-type-specific regulatory activity

We first assessed whether designed sequences were expected to have high regulatory activity in the target cell type and low activity in the background cell types. We used the Enformer-based activity predictor released with DNA-Diffusion, which jointly predicts activity across five human cell lines and achieved test-set Pearson correlations of 0.81–0.83 for K562, HepG2 and SK-N-SH [23, 33]. The predictor assigned each generated sequence an activity score in each evaluated cell type. For each method and target cell type, we assessed cell-type specificity by comparing predicted activity distributions between the target and background cell types (Methods 5.3.1). This comparison quantified the separation between intended target activity and potentially unwanted background activity. We also compared predicted target-cell activity distributions across methods to assess the strength of the intended regulatory activity.

The activity predictions revealed differences in both cell-type specificity and target-cell activity across methods (Fig. 2a and Supplementary Fig. S1). For K562, Simulated annealing, AdaLead, FastSeqProp and TACO showed the clearest separation between predicted target and background activities. RegLM and DNA-Diffusion also generated sequences with a predicted preference for K562, but the separation was generally smaller. For HepG2, Simulated annealing, AdaLead, FastSeqProp and RegLM produced sequences with higher predicted activity in HepG2 than in either background cell type. These four methods also produced elevated predicted SK-N-SH activity in sequences designed for SK-N-SH. The two D3 variants, D3 Transformer and D3 Convolutional, showed limited separation between target and background activity distributions. TACO produced elevated predicted activity in some settings, but substantial overlap between target and background distributions indicated weaker cell-type specificity. Thus, high predicted target-cell activity was not consistently accompanied by clear separation from background activity, and the extent of this separation varied with both the generation method and the target cell type.

**Fig. 2:**
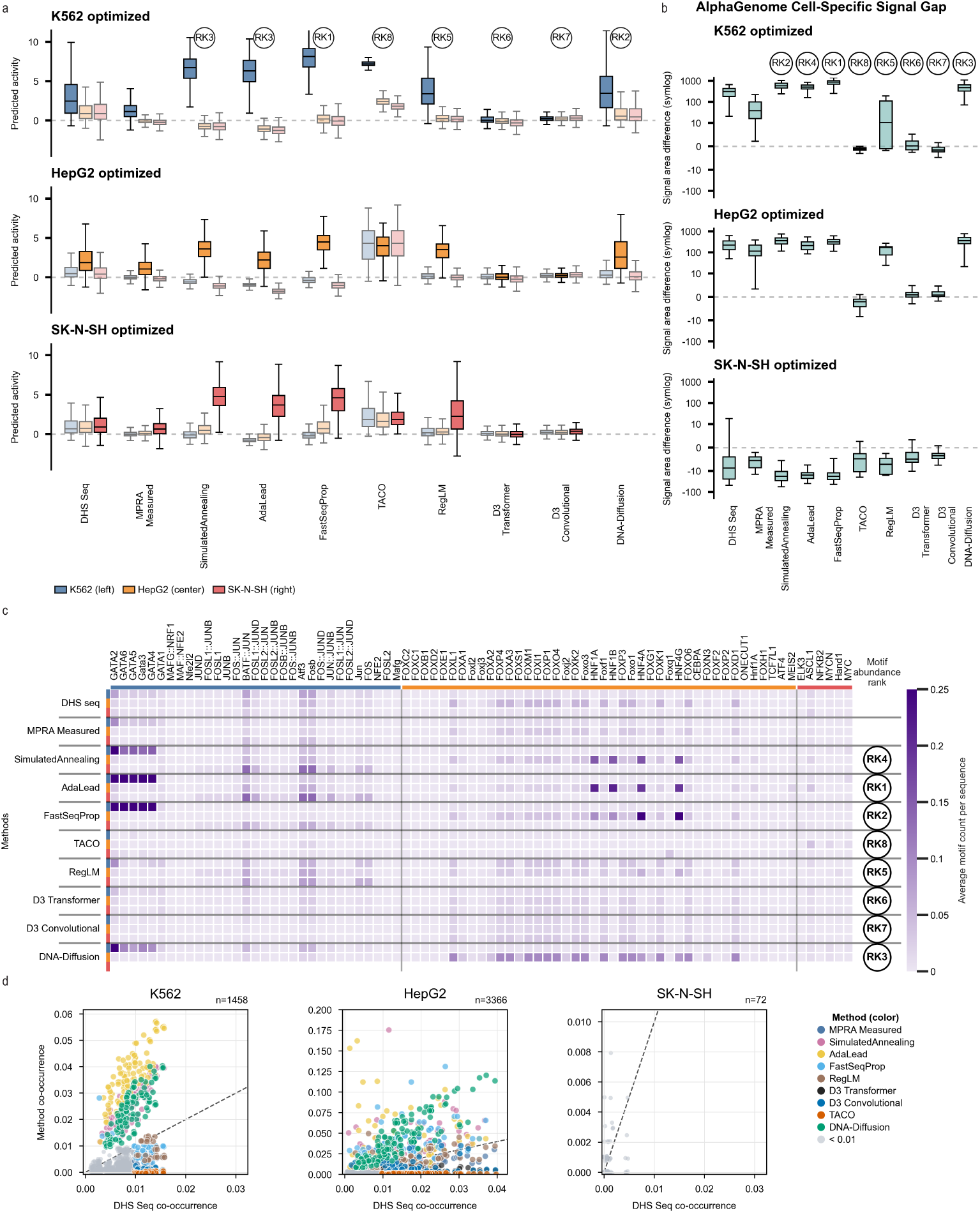
Predicted functionality and biological plausibility of reference and generated regulatory DNA. **a,** Predicted activity scores from the released Enformer-based MPRA predictor. Blue, orange and red boxes show K562, HepG2 and SK-N-SH; darker boxes identify the target. References are DHS sequences and experimentally measured, cell-specific MPRA sequences (Methods 5.1). Boxes show medians and interquartile ranges; whiskers extend to the most extreme values within 1.5 interquartile ranges. Outliers are omitted. RK labels rank generators by mean MinGap (Supplementary Fig. S1a), averaged equally across targets. DNA-Diffusion’s absent SK-N-SH mean MinGap is replaced by the arithmetic mean of the other seven generators’ SK-N-SH mean MinGap values for ranking only. **b,** AlphaGenome DNase area gaps. For each available library, up to 50 sequences with the largest activity-predictor MinGap were selected for AlphaGenome scoring. The DNase area gap was defined as the target-cell signal area minus the larger of the two non-target-cell signal areas. Areas were calculated by trapezoidal integration of unsmoothed DNase profiles over the original sequence coordinates, in signal *×* bp units. Each box summarizes sequence-level DNase area gaps. Boxes show medians and interquartile ranges; whiskers extend to the most extreme values within 1.5 interquartile ranges. The axis uses a symmetric logarithmic scale. RK labels show the chromatin-accessibility metric ranks in Fig. 1e. **c,** Mean motif-occurrence counts per sequence. Within each dataset, rows follow K562, HepG2 and SK-N-SH. Columns show the selected cell-associated motif blocks. Values at or above 0.25 share the maximum colour. **d,** Motif-pair frequencies in MPRA reference and generated libraries plotted against DHS reference frequencies. A motif pair requires non-overlapping PWM hits within the same sequence. Each point represents a comparison-dataset–pair observation; comparisons with both frequencies zero are omitted. The displayed *n* counts these observations before the HepG2 cap. Grey points satisfy max(*F*_DHS_*, F*_comparison_) *<* 0.01 and remain in the analysis. The dashed identity line indicates equal frequencies. The HepG2 y axis stops at 0.2, while all 11 frequencies above this limit remain in Supplementary Table S1. Empty DNA-Diffusion/SK-N-SH values denote unavailable generation.

We also included reference sequences corresponding to the two types of training data described above (Fig. 2a and Supplementary Fig. S1), with the MPRA reference set restricted to sequences with high experimentally measured activity in the target cell type and low activity in the background cell types (Methods 5.1). Both reference sets were scored using the same activity predictor as the generated sequences, allowing design performance to be compared with that of existing regulatory sequences.

### 2.3 Biological plausibility

We next examined additional regulatory properties to provide complementary evidence of biological plausibility and potential functionality. We first considered chromatin accessibility, which influences whether transcription factors can bind regulatory DNA and regulate gene expression [2, 3]. We used AlphaGenome [34] to predict DNase signals for up to 50 candidates per sequence set, preselected by the released activity predictor’s MinGap (Supplementary Fig. S1a). The signal gap was the target-cell DNase area minus the largest non-target-cell area (Methods 5.3.3). This analysis used a separate prediction model, but candidate selection depended on the activity evaluator. Positive gaps therefore indicate higher predicted accessibility in the target cell type.

These analyses supported cell-type-selective accessibility for several methods in K562 and HepG2 among the preselected candidates (Fig. 2b and Supplementary Fig. S2). For K562, the DHS seq and MPRA-measured reference sets showed positive signal gaps, as did sequences generated by Simulated annealing, AdaLead, FastSeqProp, RegLM and DNA-Diffusion. In comparison, D3 Transformer, D3 Convolutional and TACO produced gaps close to zero or below zero, providing little support for preferential accessibility in K562. A similar pattern was observed for HepG2.

For SK-N-SH, however, most sequence sets showed negative signal gaps, including both reference sets and generated sequences. Thus, the predicted accessibility signals did not support the target-cell preference observed in the activity-based assessment for several methods. The same discrepancy in the reference sequences suggests that limitations associated with the cellular context or accessibility predictor may contribute to the SK-N-SH results, beyond differences among generation methods.

We then examined cell-type-associated transcription factor motifs to assess whether generated sequences contained regulatory features characteristic of the intended cellular context. We quantified the average number of cell-type-associated motif occurrences per sequence and the frequency with which pairs of motifs had non-overlapping PWM matches within the same sequence (Methods 5.3.5; Fig. 2c,d and Supplementary Table S1).

Motif profiles differed substantially across methods. For K562, DNA-Diffusion, Simulated annealing, AdaLead and FastSeqProp generated sequences with high counts of GATA-family motifs (Fig. 2c). DNA-Diffusion, Simulated annealing and AdaLead also showed elevated motif-pair co-occurrence relative to the DHS seq reference (Fig. 2d). For HepG2, Simulated annealing, AdaLead and FastSeqProp concentrated motif occurrences within a limited subset of FOX- and HNF-family motifs, whereas DNA-Diffusion showed elevated counts across a broader range of HepG2-associated motifs (Fig. 2c). These methods also exhibited increased motif-pair co-occurrence relative to the DHS reference, although this pattern was less consistent across motif pairs than in K562 (Fig. 2c-d). RegLM generally showed lower and more dispersed motif counts, with more modest co-occurrence frequencies. Both D3 variants and TACO showed low occurrence and co-occurrence frequencies for most evaluated motifs.

For SK-N-SH, few cell-type-associated motifs were available for analysis, and generated sequences showed no consistent increase in their occurrence relative to the reference sets. Sparse motif signals and weak AlphaGenome-predicted accessibility preference were also observed in the reference sequences, despite their experimentally measured SK-N-SH-specific activity. This discrepancy suggests that the selected motif panel and accessibility predictions may not fully reflect the regulatory features supporting SK-N-SH-specific activity, limiting their interpretation as indicators of sequence quality in this cellular context.

Together, these analyses revealed that high predicted target-cell activity was not consistently accompanied by strong cell-type-associated motif signals. For example, TACO produced high predicted K562 activity with weak K562-associated motif signals, while RegLM showed less pronounced motif signals than several other methods with high predicted activity. The modest motif frequencies in the reference sequences also showed that experimentally supported regulatory activity need not involve marked enrichment of the selected motifs. Motif analysis therefore helped characterize the sequence features accompanying predicted activity, instead of providing a direct measure of function.

### 2.4 Sequence diversity and distributional characteristics

We next examined whether each method generated diverse candidate sequences and how these sequences related to experimentally assayed CREs. We used representations extracted from the activity predictor to place generated and MPRA reference sequences in a shared feature space, visualized using UMAP [35] (Fig. 3a; Methods 5.3.6, 5.3.7). Two variants of the local inverse Simpson’s index (LISI) [36] were used to quantify their distributions: Integration LISI (iLISI) measured local mixing between generated and reference sequences within each target cell type, with higher values indicating greater overlap; cell-type LISI among generated sequences (*cLISI_M_*) measured mixing across target cell types, with lower values indicating stronger separation (Methods 5.3.8).

**Fig. 3:**
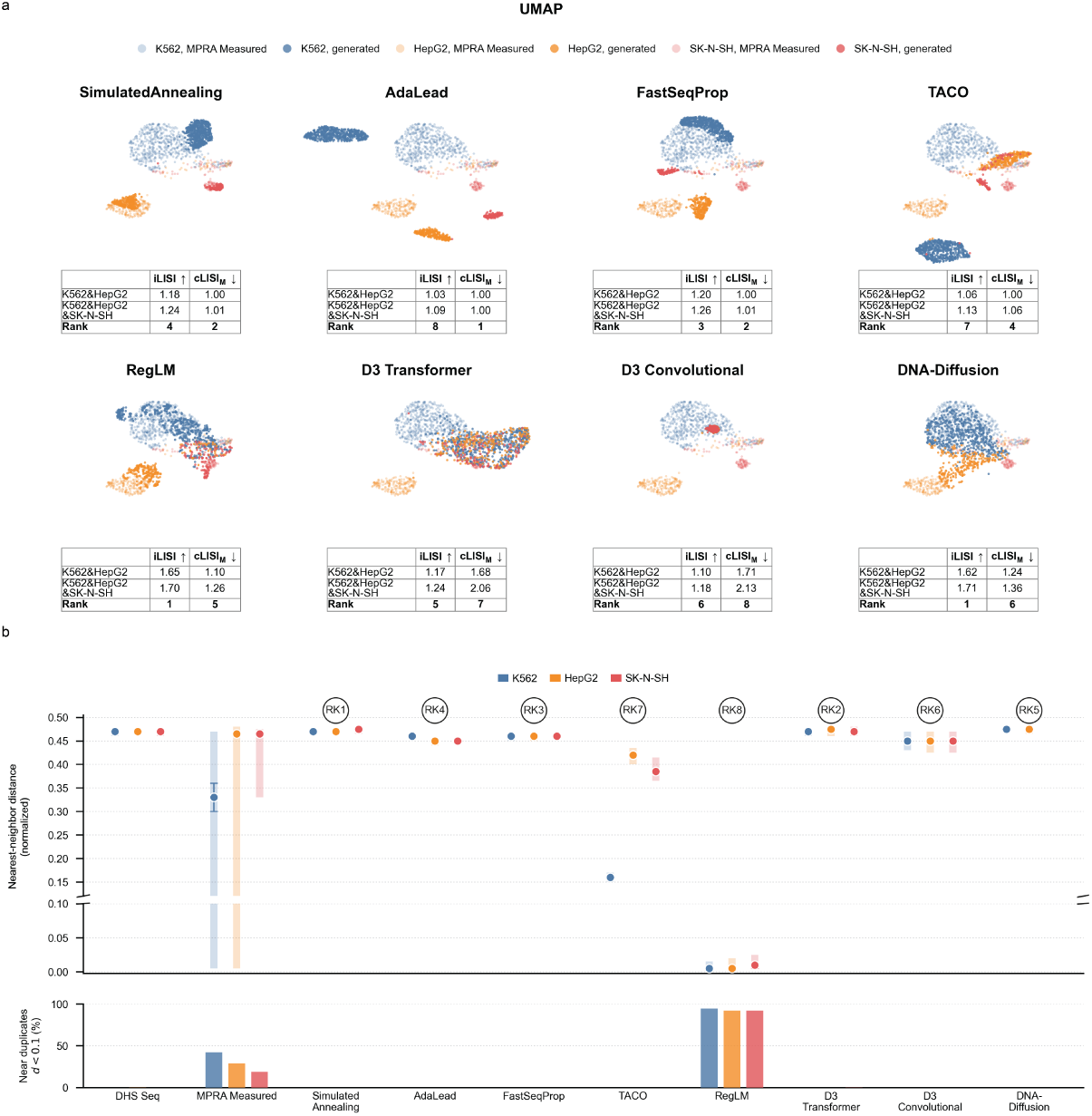
Distributional coverage, sequence diversity and cell-type specificity of generated regulatory DNA sequences. **a,** UMAP projections of MPRA-measured reference sequences (light colours) and sequences generated by eight design methods (dark colours) for K562, HepG2 and SK-N-SH. The local inverse Simpson’s index (LISI) was calculated on a Euclidean *k*-nearest-neighbour (kNN) graph with *k* = 30. Integration LISI (iLISI) quantifies local mixing between MPRA-measured and generated sequences, with higher values indicating greater distributional overlap. Cell-type LISI (cLISI) quantifies mixing among cell types, with lower values indicating stronger cell-type separation. cLISI_M_ denotes cLISI calculated among sequences generated by each method, whereas cLISI_N_ denotes cLISI calculated among measured MPRA sequences used as the common reference. The common reference values of cLISI_N_ were 1.10 for the K562–HepG2 comparison and 1.25 for the K562–HepG2–SK-N-SH comparison. Inset tables report LISI values for the K562–HepG2 and K562–HepG2–SK-N-SH comparisons. The three-cell-type values for DNA-Diffusion are reference-factor-corrected estimates because this method generated sequences only for K562 and HepG2. Each method-specific UMAP was fitted independently, aligned to the shared MPRA projection and displayed with fixed reference coordinates. LISI was calculated before projection. Inset rank rows correspond to the overview metric ranks. **b,** Normalized nearest-neighbour edit distance within each cell type for DNase I hypersensitive-site (DHS) sequences, MPRA-measured sequences and sequences generated by the eight methods (top). Dots show medians, and pale thick lines indicate the interquartile range (25th–75th percentiles). Dark thin error bars indicate 95% percentile bootstrap confidence intervals for the median. These intervals were estimated from 1,000 resamples with replacement from finite per-sequence distances within each dataset–cell-type group. Higher values indicate greater within-cell-type sequence diversity. The bottom panel shows the percentage of sequences with a normalized nearest-neighbour edit distance strictly below 0.1. Lower percentages indicate less sequence redundancy within the evaluated set.

The methods exhibited distinct combinations of distributional overlap and cell-type separation. In the K562–HepG2 comparison shared by all eight methods, RegLM and DNA-Diffusion showed relatively high overlap with the MPRA reference distribution (iLISI values), and also relatively high mixing among sequences designed for different target cell types (*cLISI_M_* values). Simulated annealing, AdaLead and FastSeqProp showed less reference overlap (iLISI), while maintaining stronger separation between target cell types (*cLISI_M_*). The three-cell-type comparisons showed the same broad patterns for the seven methods supporting SK-N-SH-targeted generation. The coverage-adjusted three-cell-type values for DNA-Diffusion are provided only as approximate references for comparison.

We then assessed nucleotide-level diversity within each library using normalized edit distances between each sequence and its closest neighbour for the same target cell type (Fig. 3b; Methods 5.3.9). Larger distances indicate more distinct candidates, whereas values approaching zero identify sequences that differ from a neighbour by very few nucleotide edits. To highlight this near-zero region, we also quantified the percentage of near duplicates (normalized distance *<* 0.1) within each library. A higher percentage indicates that a larger fraction of generated sequences have a highly similar neighbour. Most methods produced normalized nearest-neighbour edit distances of approximately 0.44–0.48, comparable to those of the DHS reference sequences, with no near duplicates detected. RegLM generated the least diverse libraries, with nearest-neighbour distances of approximately 0.01 and near-duplicate percentages exceeding 90% across all three cell types.

Viewed alongside the activity and motif analyses, these results show that limited overlap with the MPRA reference distribution can coexist with strong predicted regulatory performance, as illustrated by FastSeqProp. TACO’s stronger separation between target-cell libraries relative to RegLM did not translate into consistently stronger predicted activity specificity. Both D3 variants also generated diverse nucleotide sequences without clear cell-type separation in either feature space or predicted activity. These comparisons suggest that the value of exploring regulatory sequence space depends on producing distinct candidates that retain the desired regulatory properties. Neither proximity to the reference distribution nor sequence diversity alone establishes the usefulness of a generated library.

### 2.5 Computational efficiency

To complement the assessment of sequence quality, we evaluated the computational resources required to generate candidate libraries. We measured generation throughput and peak total memory usage across five runs on a single NVIDIA A100 GPU, using each method’s recommended generation settings and the filtering protocol (Methods 5.3.10). D3 Convolutional and TACO achieved the highest throughputs while requiring the least memory.

Mean peak total memory usage ranged from 1,222 to 8,021 MiB (Fig. 4a). Generation throughput also differed markedly among methods (Fig. 4b). D3 Convolutional achieved the highest mean throughput at 34.77 sequences per second, approximately 434.6 times that of AdaLead at 0.08 sequences per second.

**Fig. 4:**
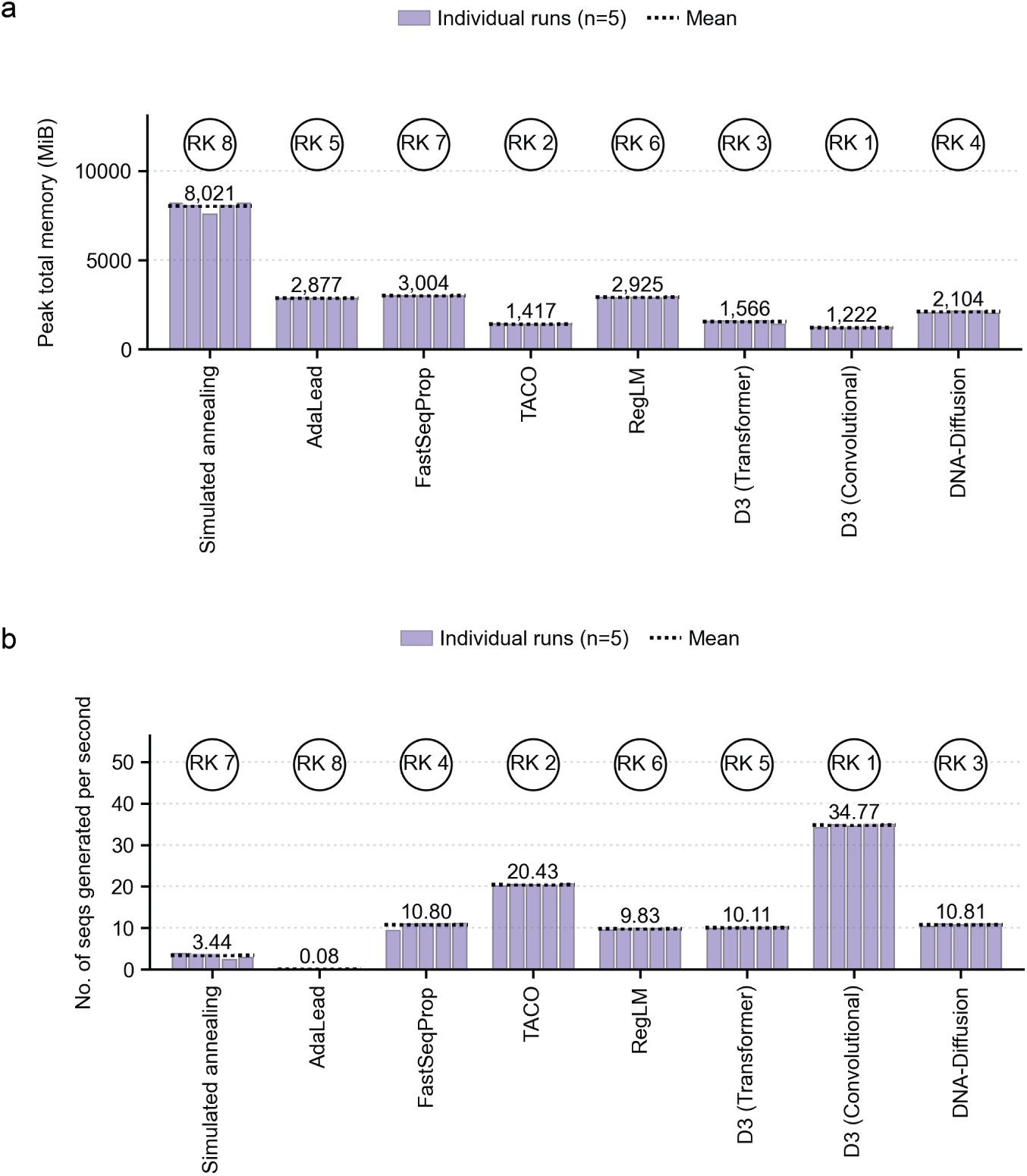
Peak memory usage and generation throughput. Eight methods were evaluated over five generation runs on one NVIDIA A100 GPU under the reported energy-threshold setting of *−*0.5 (Methods 5.3.10). Training and reinforcement-learning fine-tuning costs were excluded. **a,** Maximum combined CPU and GPU memory during generation, in MiB. **b,** Generated sequences per second. Purple bars show individual runs (*n* = 5 per method); dotted lines and annotations show arithmetic means. RK labels favour lower memory and higher throughput. Comparisons used supported target conditions; DNA-Diffusion did not generate an SK-N-SH library.

These throughput differences may partly reflect the computational procedures used during generation. D3 samples from a trained diffusion model, while TACO generates sequences from a policy previously optimized through reinforcement learning; neither requires a separate predictor-guided optimization loop for each candidate. Simulated annealing, AdaLead and FastSeqProp instead perform repeated activity evaluations during sequence optimization, adding computational work during generation. Nevertheless, FastSeqProp’s gradient-based procedure achieved throughput comparable to several direct-generation methods, indicating that iterative optimization does not necessarily entail slow generation.

## 3 Discussion

RegDesign-Bench provides a systematic comparison of eight cell-type-specific CRE design methods across predicted regulatory performance, biological plausibility, sequence diversity and computational efficiency. FastSeqProp and DNA-Diffusion performed strongly across functionality and biological-plausibility measurements, while other methods showed advantages in sequence diversity or generation efficiency. These profiles support method selection based first on the desired regulatory activity and specificity, with candidate diversity and computational requirements considered in relation to the intended application.

Differences in how functional objectives guide design may help explain these performance patterns. Simulated annealing, AdaLead and FastSeqProp repeatedly optimize candidates against activity predictions, directly favouring sequence changes that improve the specified target–background contrast. Their concentrated motif profiles are consistent with preferential amplification of regulatory features rewarded by this objective. Direct-generation methods depend on how effectively learned conditioning translates into the desired properties during sampling. The weak cell-type separation of the evaluated D3 libraries suggests that their conditioning did not produce clearly differentiated regulatory profiles, whereas DNA-Diffusion showed stronger cell-type-associated signals. Because we retained each implementation’s original training setup and generation settings, these differences cannot be attributed solely to architecture or design strategy. Training data, conditioning objectives and sampling procedures remain plausible contributors, whose effects would require controlled comparisons to resolve.

The biological analyses also identify limits to interpreting individual regulatory readouts. High predicted activity was not consistently accompanied by strong signals for the selected cell-type-associated motifs, while elevated motif occurrence and co-occurrence did not necessarily resemble reference patterns. Marked motif enrichment was also absent from some experimentally supported MPRA reference sequences, indicating that enrichment of the selected motifs is not a prerequisite for regulatory activity. This distinction was particularly apparent in SK-N-SH, where weak accessibility preference and sparse motif signals occurred in reference sequences as well as generated libraries. The shared discrepancy suggests that the selected motif panel and accessibility predictions may incompletely capture relevant regulatory features, limiting their ability to distinguish generation failures from evaluation limitations in this cellular context.

Reference similarity and candidate diversity likewise require separate interpretation. RegLM and DNA-Diffusion both showed relatively high overlap with the MPRA reference distribution, yet their libraries differed substantially in nucleotide-level redundancy. Reference overlap therefore provides little assurance that a method produces many distinct candidates. Conversely, FastSeqProp generated diverse sequences with strong predicted regulatory properties despite limited reference overlap. Together, these observations argue against treating reference similarity as a sufficient measure of library quality.

The methods included here represent a practical subset of regulatory DNA design approaches, selected for their distinct strategies and executable workflows supporting the evaluated cellular contexts. Other relevant approaches involve different biological settings or design constraints. For example, Taskiran et al. demonstrated enhancer design in fly brain cell types and human melanoma states [15], while Frömel et al. developed motif-based design strategies for specificity among hematopoietic progenitor states [21]. Applying these workflows here would require adapting their cell-specific predictors or regulatory rules. DeepSEED generates flanking sequences around predefined regulatory elements for constitutive or inducible promoter design [37]; its inclusion would require specifying these fixed elements and adapting the design objective to the evaluated target–background comparisons. These adaptations introduce choices beyond the released configurations assessed here. Future inclusion of such approaches would benefit from explicitly standardized adaptation procedures and design constraints.

A major limitation is the aggregate ranking that introduces dependencies on metric selection and weighting. Summing metric ranks gives greater influence to dimensions containing more metrics, while correlated measurements may reinforce similar aspects of performance. Individual results should therefore accompany overall rankings when selecting methods for a particular application. Unequal target-cell coverage is especially relevant to DNA-Diffusion. The observed K562–HepG2 results provide the firmer basis for comparison across all eight methods.

Also, in this study, the generated sequences were evaluated computationally. Prospective MPRA measurements across matched target and background cell types would connect these computational profiles to experimental design success. Sampling candidates across activity scores, motif profiles and degrees of reference overlap would help determine which measurements are informative for experimental selection. A useful endpoint would be the proportion of nonredundant candidates meeting prespecified target-activity and background-activity thresholds. This would combine regulatory performance and sequence diversity into a concrete measure of how many distinct, experimentally supported CREs a method provides for the intended application.

## 4 Conclusion

We present a systematic, multidimensional benchmark of eight methods for cell-type-specific regulatory DNA design. The framework evaluates functionality and biological plausibility, sequence diversity and computational efficiency through complementary measurements. No single generation strategy was uniformly optimal, and method selection should reflect the intended design objective and cellular context. FastSeqProp ranked first overall and combined strong predicted activity specificity with nonredundant sequence generation. DNA-Diffusion ranked second under coverage-adjusted aggregation and combined high MPRA-reference overlap with low redundancy in its two supported cell types. D3 Convolutional offered the highest throughput and lowest peak memory usage when generation efficiency was the primary constraint. These recommendations remain specific to the tested datasets, cell lines, predictors and metric definitions. Evidence was weaker for SK-N-SH, and DNA-Diffusion provided no SK-N-SH-targeted designs. Because generated-sequence function was assessed computationally, prospective MPRA validation across matched target and non-target cells remains necessary. The benchmark exposes these trade-offs and boundaries to support method selection and assessment of future generators across broader cell types, assays and design objectives.

## 5 Methods

### 5.1 Data preparation

The benchmark used published experimental datasets as references; no new wet-laboratory experiment was performed in this study. The MPRA data and DHS reference sequences were obtained from the public resources associated with Malinois and DNA-Diffusion [17, 23].

The MPRA table contained 798,064 sequences with experimentally measured activity in K562, HepG2 and SK-N-SH, expressed as log_2_ fold change. Cell-specific references required target-cell log_2_ FC *>* 1.0 and log_2_ FC *<* 0.0 in both non-target cells. Filtering retained 3,813 K562, 1,587 HepG2 and 1,208 SK-N-SH sequences. We retained all 4,000 DHS reference sequences per cell line. Their selection is described in Malinois Supplementary Table 12 [17].

### 5.2 Method and generation hyperparameter setup

We followed each method’s documented generation workflow and retained its released checkpoint and recommended settings unless specified below. Each benchmark library contained 1,000 sequences for a supported target cell line.

#### 5.2.1 Malinois-guided optimization

Malinois is the predictive model used to guide three optimization algorithms: Simulated annealing, AdaLead and FastSeqProp [17, 27–29]. We used the generation routine in `boda2/src/generate.py`. The `--target_feature` argument was set to 0, 1 or 2 for K562, HepG2 or SK-N-SH, respectively. Each algorithm generated 1,000 sequences per target cell line.

#### 5.2.2 RegLM

RegLM uses autoregressive generation conditioned on activity labels and was trained on MPRA data [30]. We used the released human-enhancer checkpoint from https://zenodo.org/records/10669334. The target-cell input label was 3 and each non-target-cell label was 0. We generated 1,000 sequences per target for K562, HepG2 and SK-N-SH.

#### 5.2.3 DNA Discrete Diffusion (D3)

D3 uses discrete diffusion for conditional DNA generation [31]. We evaluated the released v1 MPRA checkpoints with convolutional and transformer denoisers as separate methods. Other D3 tasks and later versions were outside this comparison. Target-cell activity was set to 2.0 and non-target-cell activities to *−*2.0. Sampling requested 1,000 sequences per target cell type with a batch size of 256 and stopped once the requested total was reached. The evaluated libraries contained 1,000 sequences per target.

#### 5.2.4 DNA-Diffusion

DNA-Diffusion uses a diffusion model trained on DHS sequences [23]. We generated sequences for K562 and HepG2, the benchmark targets supported by its released check-point; no SK-N-SH library was available. We set `sampling.guidance_scale=1.0, sampling.sample_batch_size=10` and `sampling.number_of_samples=1000`.

#### 5.2.5 TACO

TACO is a reinforcement-learning method for generating yeast and human promoter sequences, with RegLM serving as its base generative model [30, 32]. Following the official tutorial, we configured `reinforce_mbo.sh` with `tasks=("k562", "hepg2", "sknsh"), batch size=200`, and `max iter=5`. The original script first performs reinforcement learning and then conducts inference using the resulting checkpoint. To enable direct inference from a trained checkpoint without repeating the reinforcement-learning stage, we implemented a separate inference routine. The TFBS reward was set to 0.1, while all other settings were kept consistent with the official tutorial.

### 5.3 Evaluation metrics

#### 5.3.1 Activity prediction and MinGap scoring

We used the Enformer-based MPRA predictor released with DNA-Diffusion [23, 33] for regulatory sequence activity prediction.

For sequence *i* designed for target cell *c*, activity specificity was summarized by

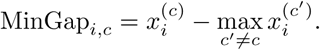

Rows with a non-finite score in any evaluated cell were excluded. We calculated the mean MinGap within each method–target library, then averaged equally across the three target-cell means for activity ranking. Reference datasets were excluded from method ranks. DNA-Diffusion’s missing SK-N-SH mean was estimated as described in Methods 5.4.2.

#### 5.3.2 Signed Wasserstein-1 score

For each generation method and target cell type *c*, we quantified cell-type specificity by comparing the predicted target activity 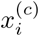 with the strongest off-target activity for each sequence *i*. The latter was defined as 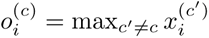, where the maximum was taken across the two non-target cell types. Let *µ_c_*and *ν_c_* denote the corresponding empirical distributions of target and off-target activities, respectively.

We computed the standard one-dimensional Wasserstein-1 distance between *µ_c_* and *ν_c_* using its empirical quantile-function representation [38, 39]. Because this distance is non-negative and does not encode the direction of the distributional shift, we defined an analysis-specific signed Wasserstein-1 score as

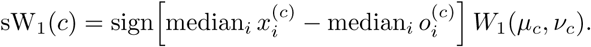

When the median activities were numerically identical, the sign was determined from the corresponding difference in mean activity. Positive scores indicate higher predicted activity in the target cell type than in the strongest off-target cell type, whereas negative scores indicate the opposite. The absolute score quantifies the overall separation between the two activity distributions.

#### 5.3.3 AlphaGenome-based DNase signal prediction

DNase signal prediction was performed using AlphaGenome [34] for three cell lines: K562, HepG2, and SK-N-SH. For each method and target cell, DNA sequences were paired with the released activity predictor’s score matrix of shape (*N,* 3), with columns ordered as K562, HepG2, and SK-N-SH. Cell specificity was ranked using a min-gap score,

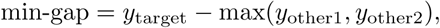

where *y* denotes predicted regulatory activity. Rows containing non-finite values in any of the three cell lines were excluded, and no additional threshold on target or off-target predictions was applied. Sequences were selected in descending MinGap order, using a requested selection size of 50 per available reference or generated library. This preselection was applied to both reference datasets and generated libraries.

For AlphaGenome inference, the input length was the smallest supported length in the installed Python client that could accommodate the original sequence. Each selected sequence was cleaned and center-padded with “N” bases to this length. The padded sequence was submitted to `predict_sequence` with `OutputType.DNASE` [34]. Ontology terms were `EFO:0002067, EFO:0001187` and `EFO:0003072` for K562, HepG2 and SK-N-SH, respectively. Returned tracks matching each cell line were identified from metadata and averaged when multiple tracks were available. The predicted signal was then cropped back to the original unpadded sequence coordinates. Signal area was computed from the unsmoothed DNase profile by trapezoidal integration. Bin spacing was the original sequence length divided by the number of cropped signal bins, expressed in base pairs. Areas and area gaps were reported in AlphaGenome-predicted DNase signal units *×* bp. Cell-specific enrichment was summarized as the target-cell signal area minus the larger area from the two non-target cells. Smoothing was used only to display signal profiles and did not affect area calculations or selection of the largest-gap candidate.

#### 5.3.4 Cell-type-associated transcription factor and motif curation

Cell-type-associated motif sets were derived from DNase-seq data for K562, HepG2 and SK-N-SH. Sequences from cell-type-associated accessible regions were extracted from the hg38 reference genome and scanned against the JASPAR 2026 non-redundant CORE vertebrate motif collection, using up to 20,000 sequences per cell type. The 200 highest-ranked motifs per cell type were retained as candidates.

To refine these candidate sets, we searched published studies in Nature, Cell and selected related journals for cell-type-relevant TF and motif evidence [17, 40–43]. Candidates were matched to manually curated entries for the corresponding cell type using JASPAR identifiers, TF names and aliases, and motif-family annotations. Only candidates with exact motif, motif-family or contextual TF matches to strong or partial literature entries were retained; background-only evidence was excluded. The filtered tables were then loaded for motif quantification, yielding 32, 45 and 7 unique TF/motif names for K562, HepG2 and SK-N-SH, respectively. The original PWMs were preserved for downstream analyses, with the three match categories annotated separately. Motif enrichment was interpreted as association with accessible chromatin rather than direct evidence of TF occupancy or regulatory direction.

#### 5.3.5 Transcription-factor motif quantification

Transcription-factor (TF) motif inputs were the literature-mapped selection tables provided for K562, HepG2 and SK-N-SH. Motifs were matched by identifier and name to position-weight matrices (PWMs) in the associated MEME files. Motif names were normalized without regard to letter case, and case-only name variants were merged only when they corresponded to an identical PWM. For the motif-abundance heatmap, let *C_m,c,t_* denote the total number of positive-strand exact matches to motif *t* among the *N_m,c_*sequences generated by method *m* for cell line *c*. The displayed value was the mean number of motif occurrences per sequence,

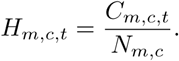

Motifs with zero counts across all method–cell-line combinations were removed, after which the first 50 non-zero motifs in the pre-ranked list for each cell line were retained. Rows represented individual method–cell-line combinations; they were arranged by method and, within each method, in the order K562, HepG2 and SK-N-SH. The colour scale was truncated at 0.25 for visualization only.

Motif co-occurrence was quantified independently using PWM-based scanning with MOODS. All motifs in the cell-specific selection files were eligible for scanning, without the top-50 restriction used for the abundance heatmap. Zero-count TF filtering was enabled using the corresponding exact-match count matrices. PWMs were converted to log-odds matrices using a uniform nucleotide background and a pseudocount of 0.01. Following the scanning configuration of Frömel *et al.*[21], motif matches were called on both DNA strands using a per-match threshold of *P* = 5 *×* 10^−5^. A motif was considered present in a sequence when at least one qualifying match was detected. A motif pair was counted only when at least one qualifying hit for each motif formed a pair of non-overlapping sequence windows. Adjacent windows were allowed, with no additional gap required. Each motif pair contributed at most one count per sequence. Thus, for motifs *a* and *b*, their sequence-level co-occurrence frequency was calculated as

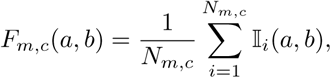

where 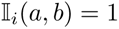 if sequence *i* contains at least one pair of non-overlapping qualifying hits for motifs *a* and *b*, and 0 otherwise. Highly similar motif pairs were excluded when their best aligned PWM Pearson correlation was at least 0.85, considering both forward and reverse-complement orientations. Co-occurrence frequencies used the number of valid sequences in each dataset as the denominator. The DHS references contained 4,000 sequences per cell; the filtered MPRA references contained 3,813, 1,587 and 1,208 sequences, respectively. No minimum-support cutoff was imposed on motif pairs; pairs observed in at least one scanned sequence were retained after the stated filters. In Fig. 2d, points with max(*F*_DHS*,c*_*, F_m,c_*) *<* 0.01 were coloured grey; low-frequency values were retained. The complete reference and generated frequency matrix is supplied in Supplementary Table S1. Its 547 pair columns form a cross-cell union. Exported zeros can represent pairs absent from a cell-specific result dictionary, whereas blank fields denote an unavailable dataset.

#### 5.3.6 Sequence embedding extraction

We extracted embeddings using the released Enformer-based MPRA predictor from DNA-Diffusion [23, 33]. Sequences were centre-cropped or padded with “N” to 200 bp and one-hot encoded over A/C/G/T; ambiguous bases were encoded as zeros. The `enformer-pytorch` backbone returned hidden representations using `return_only_embeddings=True`. The checkpoint’s learned attention-pooling layer reduced positional representations to one vector per sequence. Attention logits were normalized across positions with softmax, then used to calculate a weighted average. Each library was saved as a float32 matrix of shape *N × D*.

#### 5.3.7 UMAP visualization

We visualized generated and measured MPRA sequences using UMAP [35]. Embeddings containing non-finite values were excluded. For each cell type, the sample size was the minimum valid count across the MPRA reference and all available generated libraries. Sampling was without replacement using random generators derived from seed 42. The same sampled reference was reused across methods, with equal numbers of reference and generated sequences per cell type.

Feature-wise standardization was fitted exclusively on the pooled sampled MPRA reference. PCA was also fitted on this reference when dimensionality reduction was required, retaining at most 50 principal components. These transformations were applied unchanged to generated sequences. UMAP was fitted to the reference alone and independently to each combined reference–method dataset. Fits used `n_components=2, n_neighbors=30, _dist=0.25, metric=euclidean` and `random_state=42`.

Method-specific projections were aligned to the reference-only projection using shared MPRA points as anchors. The similarity transformation allowed rotation or reflection, uniform scaling and translation. For display, aligned MPRA coordinates were replaced with fixed reference coordinates. Projections provided qualitative visualizations; quantitative mixing scores were calculated in the transformed embedding space before projection.

#### 5.3.8 LISI-based evaluation

We quantified local mixing in the standardized, PCA-transformed Enformer space using an unweighted local inverse Simpson’s index (LISI) [36]. Neighbourhoods used Euclidean distance with *k* = 30. For categorical label *y*, let *p_i,l_* denote the proportion of neighbours of sequence *i* assigned to category *l*. Local and aggregate scores were

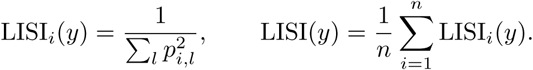

Scores were calculated before UMAP projection and alignment.

Integration LISI (iLISI) quantified mixing between measured MPRA reference and generated sequences within each target cell type [36, 44]. Source labels were N for the MPRA reference and M for generated sequences. Let *C*_2_ = {K562, HepG2} and *C*_3_ = {K562, HepG2, SK-N-SH}. For either cell-type set,

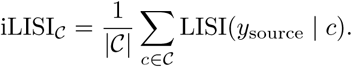

Cell-specific scores received equal weights. Both two-cell and three-cell iLISI summaries ranged from 1 to 2 because each comparison used two source categories. Higher values indicated greater local overlap with the measured MPRA reference.

Cell-type LISI quantified mixing among cell labels separately within reference and generated libraries:

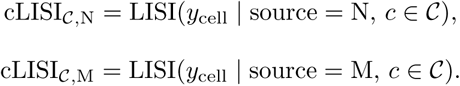

Here, *cLISI_N_* used only measured MPRA sequences and *cLISI_M_* only generated sequences. Raw cLISI ranged from 1 to 2 for *C*_2_ and from 1 to 3 for *C*_3_. Lower values indicated stronger separation among cell-type libraries. Two-cell results were directly available for all eight methods. Three-cell results were directly calculated for the seven methods supporting SK-N-SH generation. DNA-Diffusion estimates used the adjustment in Methods 5.4.2.

#### 5.3.9 Edit distance calculation and acceleration using BK-tree

To quantify sequence diversity within each generated set, we computed the nearest-neighbour edit distance for every DNA sequence within the same method–cell-line group. Each input file was parsed as one sequence per line; empty lines and comment lines were skipped, sequences were converted to uppercase, internal whitespace was removed, and records containing characters outside `A`, `C`, `G`, `T`, and `N` were excluded by validation.

For a sequence set 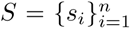, we defined the raw within-set nearest-neighbour distance of sequence *s_i_* as

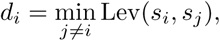

where Lev(*·, ·*) denotes the Levenshtein edit distance [45]. Exact duplicate records were treated as valid distinct neighbours and were therefore assigned *d_i_*= 0. To account for sequence-length differences, we also computed a normalized distance,

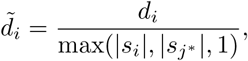

where *s_j_∗* is the nearest-neighbour sequence used for *d_i_*.

To accelerate the exact nearest-neighbour search, we built a BK-tree [46] for the unique sequences in each method–cell-line group, using Levenshtein distance as the tree metric. For non-duplicated query sequences, the identical query string was excluded from candidate matches. The BK-tree was queried with an initial radius of 10 edit operations; if no non-self neighbour was found, the radius was doubled until at least one candidate was returned or the maximum possible edit distance was reached. Because each query returns all sequences within the current radius, the minimum-distance candidate from the first non-empty search radius gives the exact nearest non-self neighbour. Computations were parallelized across query sequences using multiprocessing workers initialized with the shared BK-tree and sequence-count table. Per-group results were cached with a SHA-1 fingerprint of the input sequence set to allow reuse only when the underlying data were unchanged.

#### 5.3.10 Generation speed and memory usage

Generation throughput and peak memory usage were measured over five runs per method on one NVIDIA A100 GPU. We retained each implementation’s documented batch settings. Throughput was the number of generated sequences divided by elapsed generation time. Peak total memory was the maximum combined CPU and GPU memory consumption during generation. Arithmetic means across the five runs provided the efficiency metrics in Fig. 1e.

Libraries contained 1,000 sequences per supported target condition. DNA-Diffusion supported K562 and HepG2, whereas the other methods supported all three benchmark targets. Comparisons therefore concern generation under supported conditions rather than an identical three-target workload. The evaluated settings included the energy threshold of *−*0.5 shown in Fig. 4. Model training and reinforcement-learning fine-tuning costs were excluded.

### 5.4 Ranking and overview score calculation

#### 5.4.1 Overall Ranking

The overview in Fig. 1e was constructed from nine metrics grouped into three evaluation categories. Predicted activity, chromatin-accessibility signal, TF motif abundance, TF motif co-occurrence, iLISI, edit distance and throughput were treated as benefit metrics, for which larger values were preferred. cLISI_M_ and peak memory consumption were treated as cost metrics, for which smaller values were preferred.

Methods were first ranked separately for every metric. For a benefit metric, values were ordered from largest to smallest; for a cost metric, values were ordered from smallest to largest. We used competition ranking, such that equal values received the same rank and the next rank was skipped (for example, 1, 2, 2, 4). Let *r_ij_* denote the rank of method *i* for metric *j*. The motif metrics favour larger abundance or co-occurrence summaries as prespecified compositional proxies. The category score for category *g* was the unweighted sum of the ranks of its constituent metrics,

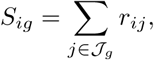

and category ranks were obtained by ranking *S_ig_* in ascending order. The grand overall score was calculated directly from all nine metric ranks,

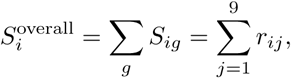

and methods were ranked in ascending order of 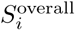, again using competition ranking for ties. Thus, a smaller category or overall score indicates better aggregate performance. No additional metric or category weights were applied; consequently, the four functionality-and-plausibility metrics, three sequence-diversity metrics and two efficiency metrics contributed four, three and two rank terms, respectively. A category or overall score was reported as unavailable if any required metric remained missing after the prespecified correction.

For visualization only, each raw metric was also min–max normalized across methods. For benefit metrics, the displayed score was

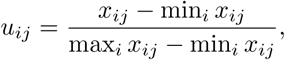

whereas for cost metrics it was

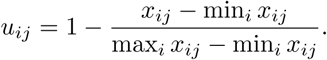

Scores were constrained to the interval [0, 1], and a metric with no between-method variation was assigned a score of 1 for all methods. Circle size in Fig. 1e represents *u_ij_*, whereas circle colour represents the metric rank. The square cells display the category and overall ranks derived from the rank sums above.

#### 5.4.2 Adjustment for unequal target-cell coverage

DNA-Diffusion generated sequences for K562 and HepG2, but not for SK-N-SH. Activity, DNase signal area gap, both motif metrics and edit distance used mean imputation for the missing target-cell score, whereas LISI summaries used a multiplicative reference-factor correction.

For activity ranking, let *A_m,c_* denote the mean MinGap in method *m*’s target-cell library *c*. Complete-coverage methods were summarized by the equally weighted mean across the three target cells. DNA-Diffusion’s missing SK-N-SH mean was estimated from the other methods:

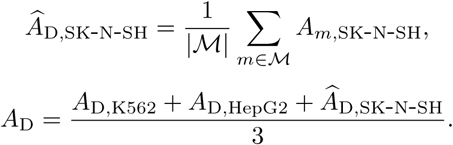

Here, *M* contains the other seven generation methods with finite SK-N-SH values; reference datasets are excluded. The same procedure was applied separately to DNase signal area gap, TF motif abundance, TF motif co-occurrence and edit distance: the missing SK-N-SH score was replaced by the arithmetic mean of the corresponding SK-N-SH scores from the other seven generation methods, excluding reference datasets.

For each metric, DNA-Diffusion’s aggregate score was then the equally weighted mean of its observed K562 and HepG2 scores and this estimated SK-N-SH score. These estimates contributed only to aggregate comparisons. No SK-N-SH-targeted DNA-Diffusion sequence or per-sequence prediction was inferred or displayed.

For Fig. 3a, LISI summaries used a multiplicative reference-factor correction. Let 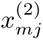 and 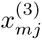 denote metric *j* under two-cell and three-cell comparisons. The correction was

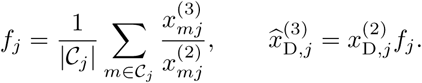

The reference set *C_j_* contained complete-coverage methods with finite values and a non-zero two-cell denominator. Factors were calculated separately for each LISI metric. Adjusted values estimated three-cell summaries without reconstructing an SK-N-SH library. Throughput and peak memory measurements were unchanged. Both adjustment procedures provide approximate comparisons and cannot establish observed SK-N-SH generation performance.

## Supplementary

**Table S1:**
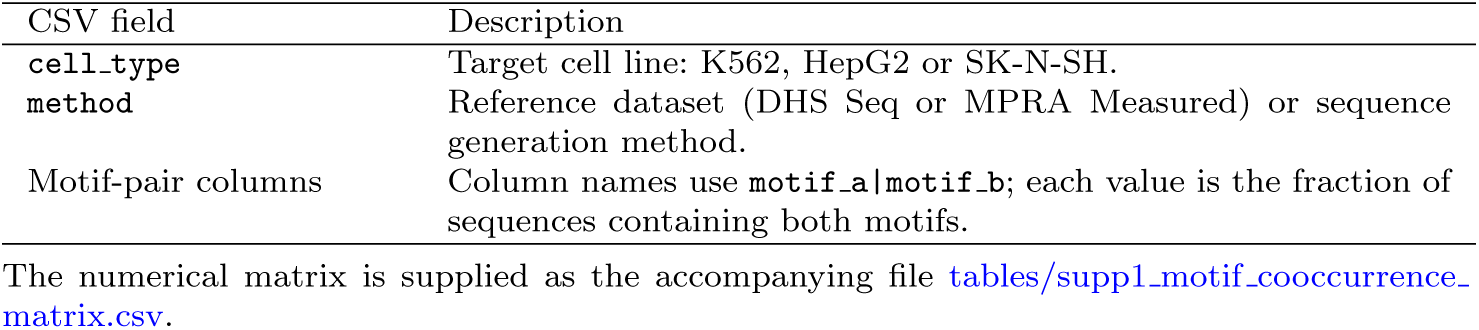
Motif-pair co-occurrence frequency matrix for reference and generated sequences. The accompanying CSV contains the frequencies used in Fig. 2d, with 30 rows representing three cell lines, two reference datasets (DHS seq and MPRA-measured sequences) and eight generation methods, and 547 motif-pair columns. Frequencies are supplied without the plotting cap, including 11 HepG2 entries above 0.2.

**Fig. S1:**
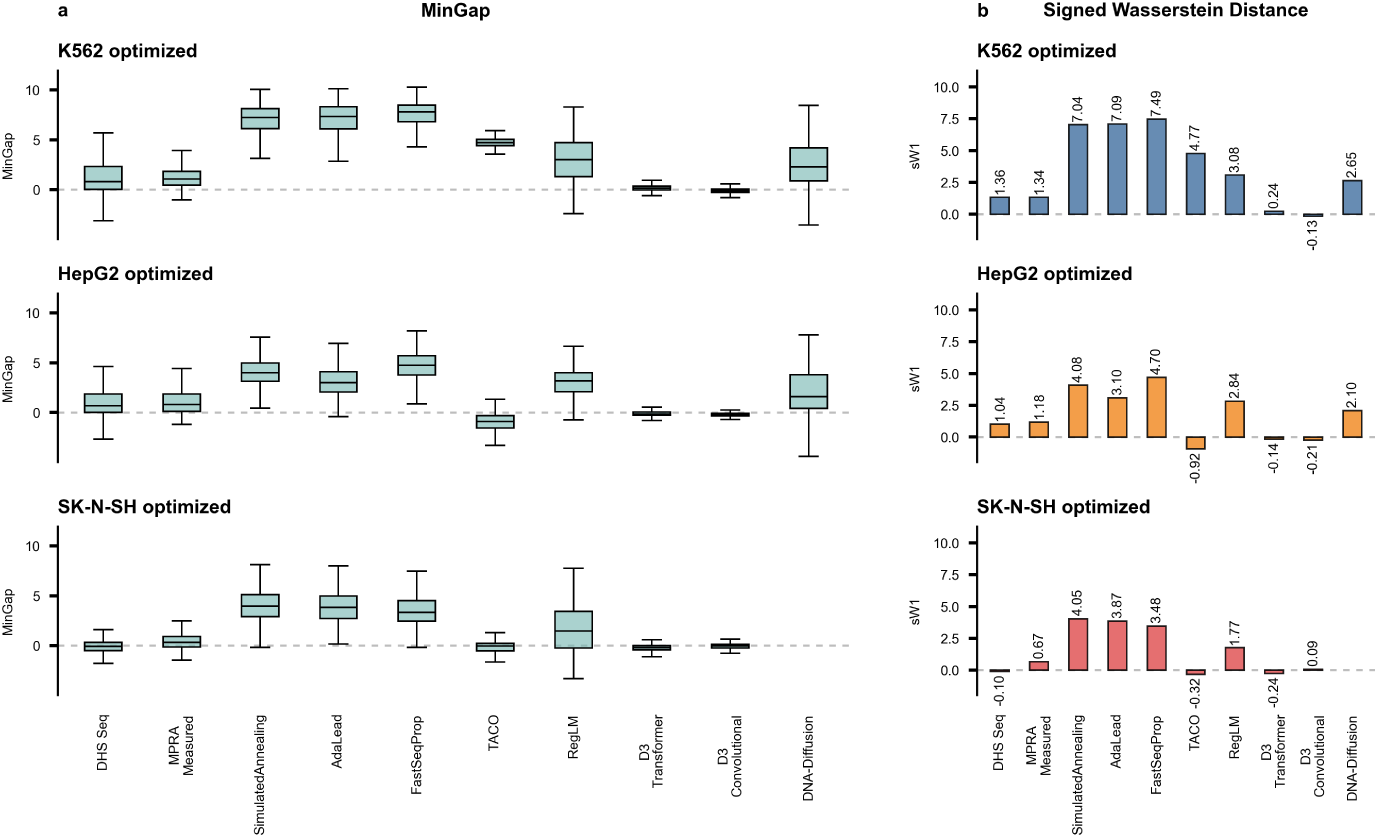
MinGap and signed Wasserstein-1 scores from the released activity predictor. Reference and generated libraries were evaluated with the Enformer-based predictor released with DNA-Diffusion [23]. **a,** Per-sequence MinGap, defined as target-cell predicted activity minus the maximum of the two non-target-cell activities. Positive values indicate target preference. Boxes show medians and interquartile ranges; whiskers extend to the most extreme values within 1.5 interquartile ranges, with outliers omitted. **b,** Signed Wasserstein-1 scores compare target and strongest off-target activity distributions. Magnitude measures distributional separation; the sign follows the median difference, with mean differences resolving numerical median ties. Annotated values report scores. Bar colours indicate K562, HepG2 and SK-N-SH in blue, orange and red, respectively. Dashed lines indicate zero. DNA-Diffusion/SK-N-SH is unavailable. These signed scores complement the mean-MinGap activity ranks.

**Fig. S2:**
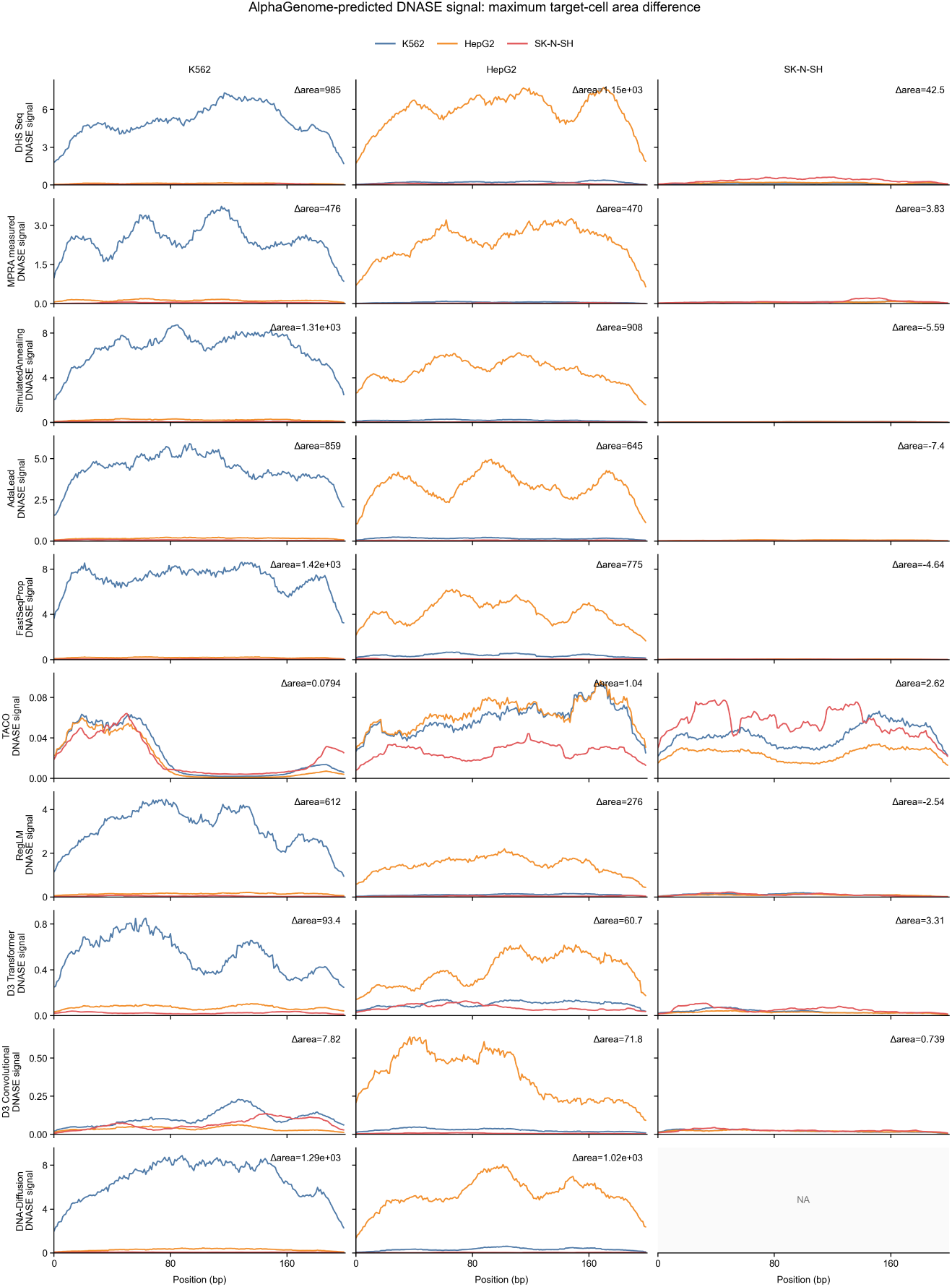
Selected AlphaGenome-predicted DNase profiles. Columns denote K562, HepG2 and SK-N-SH target libraries; rows denote reference datasets and generation methods. Curves show predicted DNase signals along the 200-bp sequence for K562 (blue), HepG2 (orange) and SK-N-SH (red). Each example has the largest DNase area gap among up to 50 candidates preselected by the released activity predictor’s MinGap. Thus panels illustrate selected examples. Annotated Δarea is the target-cell area minus the larger non-target-cell area, calculated from unsmoothed profiles. Panel-specific y-axis scales limit visual comparison of peak heights. NA denotes unavailable DNA-Diffusion/SK-N-SH generation.

## Supplementary information

Supplementary Figures S1–S3 and Supplementary Table S1 accompany this article. The complete numerical Table S1 is supplied as `supp1_motif_cooccurrence_matrix.csv`.

## Data availability

The published MPRA and DHS source datasets are described in Methods 5.1 and their original studies [17, 23]. The motif-pair frequency matrix used in Fig. 2d accompanies this manuscript as Supplementary Table S1.

## Notes

### Competing Interest Statement

The authors have declared no competing interest.

## References

[1] Wittkopp, P.J., Kalay, G.: Cis-regulatory elements: molecular mechanisms and evolutionary processes underlying divergence. Nature Reviews Genetics 13, 59–69 (2012) 10.1038/nrg3095

[2] Heinz, S., Romanoski, C.E., Benner, C., Glass, C.K.: The selection and function of cell type-specific enhancers. Nature Reviews Molecular Cell Biology 16(3), 144–154 (2015) 10.1038/nrm3949

[3] Shlyueva, D., Stampfel, G., Stark, A.: Transcriptional enhancers: from properties to genome-wide predictions. Nature Reviews Genetics 15(4), 272–286 (2014) 10.1038/nrg3682

[4] Heintzman, N.D., Hon, G.C., Hawkins, R.D., Kheradpour, P., Stark, A., Harp, L.F., Ye, Z., Lee, L.K., Stuart, R.K., Ching, C.W., Ching, K.A., Antosiewicz-Bourget, J.E., Liu, H., Zhang, X., Green, R.D., Lobanenkov, V.V., Stewart, R., Thomson, J.A., Crawford, G.E., Kellis, M., Ren, B.: Histone modifications at human enhancers reflect global cell-type-specific gene expression. Nature 459(7243), 108–112 (2009) 10.1038/nature07829

[5] Agarwal, V., Inoue, F., Schubach, M., Penzar, D., Martin, B.K., Dash, P.M., Keukeleire, P., Zhang, Z., Sohota, A., Zhao, J., Georgakopoulos-Soares, I., Noble, W.S., Yardımcı, G.G., Kulakovskiy, I.V., Kircher, M., Shendure, J., Ahituv, N.: Massively parallel characterization of transcriptional regulatory elements. Nature 639(8054), 411–420 (2025) 10.1038/s41586-024-08430-9

[6] Graybuck, L.T., Daigle, T.L., Sedeño-Cortés, A.E., Walker, M., Kalmbach, B., Lenz, G.H., Morin, E., Nguyen, T.N., Garren, E., Bendrick, J.L., Kim, T.K., Zhou, T., Mortrud, M., Yao, S., Siverts, L.A., Larsen, R., Gore, B.B., Szelenyi, E.R., Trader, C., Balaram, P., Velthoven, C.T.J., Chiang, M., Mich, J.K., Dee, N., Goldy, J., Cetin, A.H., Smith, K., Way, S.W., Esposito, L., Yao, Z., Gradinaru, V., Sunkin, S.M., Lein, E., Levi, B.P., Ting, J.T., Zeng, H., Tasic, B.: Enhancer viruses for combinatorial cell-subclass-specific labeling. Neuron 109(9), 1449–146413 (2021) 10.1016/j.neuron.2021.03.011

[7] Dimidschstein, J., Chen, Q., Tremblay, R., Rogers, S.L., Saldi, G.-A., Guo, L., Xu, Q., Liu, R., Lu, C., Chu, J., Grimley, J.S., Krostag, A.-R., Kaykas, A., Avery, M.C., Rashid, M.S., Baek, M., Jacob, A.L., Smith, G.B., Wilson, D.E., Kosche, G., Kruglikov, I., Rusielewicz, T., Kotak, V.C., Mowery, T.M., Anderson, S.A., Callaway, E.M., Dasen, J.S., Fitzpatrick, D., Fossati, V., Long, M.A., Noggle, S., Reynolds, J.H., Sanes, D.H., Rudy, B., Feng, G., Fishell, G.: A viral strategy for targeting and manipulating interneurons across vertebrate species. Nature Neuroscience 19(12), 1743–1749 (2016) 10.1038/nn.4430

[8] Vormstein-Schneider, D., Lin, J.D., Pelkey, K.A., Chittajallu, R., Guo, B., Arias-Garcia, M.A., Allaway, K., Sakopoulos, S., Schneider, G., Stevenson, O., Vergara, J., Sharma, J., Zhang, Q., Franken, T.P., Smith, J., Ibrahim, L.A., Mastro, K.J., Sabri, E., Huang, S., Favuzzi, E., Burbridge, T., Xu, Q., Guo, L., Vogel, I., Sanchez, V., Saldi, G.A., Gorissen, B.L., Yuan, X., Zaghloul, K.A., Devinsky, O., Sabatini, B.L., Batista-Brito, R., Reynolds, J., Feng, G., Fu, Z., McBain, C.J., Fishell, G., Dimidschstein, J.: Viral manipulation of functionally distinct interneurons in mice, non-human primates and humans. Nature Neuroscience 23(12), 1629–1636 (2020) 10.1038/s41593-020-0692-9

[9] Greenshpan, Y., Sharabi, O., Ottolenghi, A., Cahana, A., Kundu, K., Yegodayev, K.M., Elkabets, M., Gazit, R., Porgador, A.: Synthetic promoters to induce immune-effectors into the tumor microenvironment. Communications Biology 4(1), 143 (2021) 10.1038/s42003-021-01664-7

[10] Nissim, L., Wu, M.-R., Pery, E., Binder-Nissim, A., Suzuki, H.I., Stupp, D., Wehrspaun, C., Tabach, Y., Sharp, P.A., Lu, T.K.: Synthetic RNA-based immunomodulatory gene circuits for cancer immunotherapy. Cell 171(5), 1138–115015 (2017) 10.1016/j.cell.2017.09.049

[11] Bonfá, G., Martino, G., Sellitto, A., Blazquez-Roman, J.L., Tedeschi, F., Rinaldi, A., Silveira, C.R.F., Piro, F., Caliendo, F., Melchiorri, L., Perna, D., Tarnai, R., Heuser-Loy, C., Mastrogiovanni, F., Vadász, T., Starkey, F., Nikolados, E., Menolascina, F., Gattinoni, L., Siciliano, V.: A modular synthetic promoter toolkit for programmable gene control in activated and chronically stimulated T cells. Nature Communications (2026) 10.1038/s41467-026-77730-7

[12] Koeber, U., Matjusaitis, M., Alfazema, N., Furlong, K., Wang, Z., White, R., Hamdan, A., Dewari, P., Morisse, G., Navarette, M., Willis, R., Wang, J., Clark, M.P., Sousa, C., Hong, H.I., Sheraz, S., Southgate, B., Cholewa-Waclaw, J., Gogolok, S., Morrison, G.M., Cancino, F.G., Robertson, F., Williams, A., Rosser, S.J., Brennan, P.M., Sieger, D., Soufi, A., Quezada, S.A., Pollard, S.M.: Synthetic super-enhancers enable precision viral immunotherapy. Nature 653(8113), 232–241 (2026) 10.1038/s41586-026-10329-6

[13] Psatha, N., Sova, P., Georgolopoulos, G., Paschoudi, K., Iwata, M., Bloom, J., Ulyanova, T., Wang, H., Kirtsou, A., Vasiloudis, N.-I., Wilken, M.S., Stamatoyannopoulos, J.A., Yannaki, E., Papayanopoulou, T., Stamatoyannopoulos, G., Vierstra, J.: Large-scale discovery of potent, compact and erythroid specific enhancers for gene therapy vectors. Nature Communications 16(1), 4325 (2025) 10.1038/s41467-025-59235-x

[14] Jüttner, J., Szabo, A., Gross-Scherf, B., Morikawa, R.K., Rompani, S.B., Hantz, P., Szikra, T., Esposti, F., Cowan, C.S., Bharioke, A., Patino-Alvarez, C.P., Keles, Ö., Kusnyerik, A., Azoulay, T., Hartl, D., Krebs, A.R., Schübeler, D., Hajdu, R.I., Lukats, A., Nemeth, J., Nagy, Z.Z., Wu, K.-C., Wu, R.-H., Xiang, L., Fang, X.-L., Jin, Z.-B., Goldblum, D., Hasler, P.W., Scholl, H.P.N., Krol, J., Roska, B.: Targeting neuronal and glial cell types with synthetic promoter AAVs in mice, non-human primates and humans. Nature Neuroscience 22(8), 1345–1356 (2019) 10.1038/s41593-019-0431-2

[15] Taskiran, I.I., Spanier, K.I., Dickmänken, H., Kempynck, N., Pančíková, A., Ekşi, E.C., Hulselmans, G., Ismail, J.N., Theunis, K., Vandepoel, R., Christiaens, V., Mauduit, D., Aerts, S.: Cell-type-directed design of synthetic enhancers. Nature 626(7997), 212–220 (2024) 10.1038/s41586-023-06936-2

[16] Almeida, B.P., Schaub, C., Pagani, M., Secchia, S., Furlong, E.E.M., Stark, A.: Targeted design of synthetic enhancers for selected tissues in the Drosophila embryo. Nature 626(7997), 207–211 (2024) 10.1038/s41586-023-06905-9

[17] Gosai, S.J., Castro, R.I., Fuentes, N., Butts, J.C., Mouri, K., Alasoadura, M., Kales, S., Nguyen, T.T.L., Noche, R.R., Rao, A.S., Joy, M.T., Sabeti, P.C., Reilly, S.K., Tewhey, R.: Machine-guided design of cell-type-targeting cis-regulatory elements. Nature 634(8036), 1211–1220 (2024) 10.1038/s41586-024-08070-z

[18] Yin, C., Castillo-Hair, S., Byeon, G.W., Bromley, P., Meuleman, W., Seelig, G.: Iterative deep learning design of human enhancers exploits condensed sequence grammar to achieve cell-type specificity. Cell Systems 16(7), 101302 (2025) 10.1016/j.cels.2025.101302

[19] Kempynck, N., De Winter, S., Blaauw, C.H., Konstantakos, V., Ekşi, E.C., Dieltiens, S., Abaffyová, D., Bercier, V., Taskiran, I.I., Hulselmans, G., Spanier, K., Christiaens, V., Van Den Bosch, L., Mahieu, L., Aerts, S.: CREsted: modeling genomic and synthetic cell-type-specific enhancers across tissues and species. Nature Methods 23(5), 946–959 (2026) 10.1038/s41592-026-03057-2

[20] Chen, S., Loubiere, V., Hollingsworth, E.W., Murakami, K., Mandlburger, N., Jacinto, S.H., Dizehchi, A., Schreiber, J., Kvon, E.Z., Stark, A.: Predictive design of tissue-specific mammalian enhancers that function in the mouse embryo. Nature Genetics 58(9), 2129–2133 (2026) 10.1038/s41588-026-02729-1

[21] Frömel, R., Rühle, J., Bernal Martinez, A., Szu-Tu, C., Pacheco Pastor, F., Martinez-Corral, R., Velten, L.: Design principles of cell-state-specific enhancers in hematopoiesis. Cell 188(12), 3202–321821 (2025) 10.1016/j.cell.2025.04.017

[22] Cornejo-Páramo, P., Zhang, X., Louis, L., Li, Z., Yang, Y., Wong, E.S.: Motif-based models accurately predict cell type-specific distal regulatory elements. Nature Communications 16(1), 10370 (2025) 10.1038/s41467-025-65362-2

[23] DaSilva, L.F., Senan, S., Kribelbauer-Swietek, J.F., Patel, Z.M., Louis, L.K., Reddy, A.J., Gabbita, S., Rosen, J.D., Nussbaum, Z., Córdova, C.M.V., Wenteler, A., Weber, N., Tunjic, T.M., Mansoldo, M., Khan, T.A., Hwang, G.-H., Gardeux, V., Humphreys, D.T., Smith, C., Bejan, M., Bromley, P., Connell, W., Deplancke, B., Love, M.I., Wong, E.S., Meuleman, W., Pinello, L.: Designing synthetic regulatory elements using the generative AI framework DNA-Diffusion. Nature Genetics 58(1), 180–194 (2026) 10.1038/s41588-025-02441-6

[24] Meuleman, W., Muratov, A., Rynes, E., Halow, J., Lee, K., Bates, D., Diegel, M., Dunn, D., Neri, F., Teodosiadis, A., Reynolds, A., Haugen, E., Nelson, J., Johnson, A., Frerker, M., Buckley, M., Sandstrom, R., Vierstra, J., Kaul, R., Stamatoyannopoulos, J.: Index and biological spectrum of human DNase I hypersensitive sites. Nature 584(7820), 244–251 (2020) 10.1038/s41586-020-2559-3

[25] Heinz, S., Benner, C., Spann, N., Bertolino, E., Lin, Y.C., Laslo, P., Cheng, J.X., Murre, C., Singh, H., Glass, C.K.: Simple combinations of lineage-determining transcription factors prime cis-regulatory elements required for macrophage and B cell identities. Molecular Cell 38(4), 576–589 (2010) 10.1016/j.molcel.2010.05.004

[26] Noack, F., Vangelisti, S., Raffl, G., Carido, M., Diwakar, J., Chong, F., Bonev, B.: Multimodal profiling of the transcriptional regulatory landscape of the developing mouse cortex identifies Neurog2 as a key epigenome remodeler. Nature Neuroscience 25(2), 154–167 (2022) 10.1038/s41593-021-01002-4

[27] Laarhoven, P.J.M., Aarts, E.H.L.: Simulated annealing. In: Laarhoven, P.J.M., Aarts, E.H.L. (eds.) Simulated Annealing: Theory and Applications. Mathematics and Its Applications, vol. 37, pp. 7–15. Springer, Dordrecht (1987). 10.1007/978-94-015-7744-1_2

[28] Sinai, S., Wang, R., Whatley, A., Slocum, S., Locane, E., Kelsic, E.D.: AdaLead: A simple and robust adaptive greedy search algorithm for sequence design. arXiv (2020). 10.48550/ARXIV.2010.02141. https://arxiv.org/abs/2010.02141

[29] Linder, J., Seelig, G.: Fast activation maximization for molecular sequence design. BMC Bioinformatics 22(1) (2021) 10.1186/s12859-021-04437-5

[30] Lal, A., Garfield, D., Biancalani, T., Eraslan, G.: Designing realistic regulatory dna with autoregressive language models. Genome Research 34(9), 1411–1420 (2024) 10.1101/gr.279142.124

[31] Sarkar, A., Duran, A., Yu, Y., Lin, D.-W., Kang, Y., Somia, N., Mantilla, P., Zhou, J., Nagai, M., Tang, Z., Hanington, K., Chang, K., Koo, P.K.: Designing DNA with tunable regulatory activity using discrete diffusion. bioRxiv (2024) 10.1101/2024.05.23.595630. Preprint, version 3

[32] Yang, Z., Su, B., Cao, C., Wen, J.-R.: Regulatory dna sequence design with reinforcement learning. In: The Thirteenth International Conference on Learning Representations (2025). https://openreview.net/forum?id=F4IMiNhim1

[33] Avsec, Ž., Agarwal, V., Visentin, D., Ledsam, J.R., Grabska-Barwinska, A., Taylor, K.R., Assael, Y., Jumper, J., Kohli, P., Kelley, D.R.: Effective gene expression prediction from sequence by integrating long-range interactions. Nature Methods 18(10), 1196–1203 (2021) 10.1038/s41592-021-01252-x

[34] Avsec, Ž., Latysheva, N., Cheng, J., Novati, G., Taylor, K.R., Ward, T., Bycroft, C., Nicolaisen, L., Arvaniti, E., Pan, J., Thomas, R., Dutordoir, V., Perino, M., De, S., Karollus, A., Gayoso, A., Sargeant, T., Mottram, A., Wong, L.H., Drotár, P., Kosiorek, A., Senior, A., Tanburn, R., Applebaum, T., Basu, S., Hassabis, D., Kohli, P.: Advancing regulatory variant effect prediction with alphagenome. Nature 649(8099), 1206–1218 (2026) 10.1038/s41586-025-10014-0

[35] McInnes, L., Healy, J., Melville, J.: UMAP: Uniform manifold approximation and projection for dimension reduction. arXiv preprint arXiv:1802.03426 (2018) 10.48550/arXiv.1802.03426 arXiv:1802.03426 [stat.ML]

[36] Korsunsky, I., Millard, N., Fan, J., Slowikowski, K., Zhang, F., Wei, K., Baglaenko, Y., Brenner, M., Loh, P.-R., Raychaudhuri, S.: Fast, sensitive and accurate integration of single-cell data with Harmony. Nature Methods 16(12), 1289–1296 (2019) 10.1038/s41592-019-0619-0

[37] Zhang, P., Wang, H., Xu, H., Wei, L., Liu, L., Hu, Z., Wang, X.: Deep flanking sequence engineering for efficient promoter design using DeepSEED. Nature Communications 14(1), 6309 (2023) 10.1038/s41467-023-41899-y

[38] Vallender, S.S.: Calculation of the Wasserstein distance between probability distributions on the line. Theory of Probability and Its Applications 18(4), 784–786 (1974) 10.1137/1118101

[39] Peyré, G., Cuturi, M.: Computational optimal transport. Foundations and Trends in Machine Learning 11(5–6), 355–607 (2019) 10.1561/2200000073

[40] Pierce, S.E., Granja, J.M., Greenleaf, W.J.: High-throughput single-cell chromatin accessibility CRISPR screens enable unbiased identification of regulatory networks in cancer. Nature Communications 12, 2969 (2021) 10.1038/s41467-021-23213-w

[41] Bravo González-Blas, C., Matetovici, I., Hillen, H., Taskiran, I.I., Vandepoel, R., Christiaens, V., Sansores-García, L., Verboven, E., Hulselmans, G., Poovathingal, S., Demeulemeester, J., Psatha, N., Mauduit, D., Halder, G., Aerts, S.: Single-cell spatial multi-omics and deep learning dissect enhancer-driven gene regulatory networks in liver zonation. Nature Cell Biology 26(1), 153–167 (2024) 10.1038/s41556-023-01316-4

[42] Thirant, C., Peltier, A., Durand, S., Kramdi, A., Louis-Brennetot, C., Pierre-Eugène, C., Gautier, M., Costa, A., Grelier, A., Zaïdi, S., Gruel, N., Jimenez, I., Lapouble, E., Pierron, G., Sitbon, D., Brisse, H.J., Gauthier, A., Fréneaux, P., Grossetête, S., Baudrin, L.G., Raynal, V., Baulande, S., Bellini, A., Bhalshankar, J., Carcaboso, A.M., Geoerger, B., Rohrer, H., Surdez, D., Boeva, V., Schleiermacher, G., Delattre, O., Janoueix-Lerosey, I.: Reversible transitions between noradrenergic and mesenchymal tumor identities define cell plasticity in neuroblastoma. Nature Communications 14, 2575 (2023) 10.1038/s41467-023-38239-5

[43] Wang, L., Tan, T.K., Durbin, A.D., Zimmerman, M.W., Abraham, B.J., Tan, S.H., Ngoc, P.C.T., Weichert-Leahey, N., Akahane, K., Lawton, L.N., Rokita, J.L., Maris, J.M., Young, R.A., Look, A.T., Sanda, T.: ASCL1 is a MYCN- and LMO1-dependent member of the adrenergic neuroblastoma core regulatory circuitry. Nature Communications 10, 5622 (2019) 10.1038/s41467-019-13515-5

[44] Luecken, M.D., Büttner, M., Chaichoompu, K., Danese, A., Interlandi, M., Mueller, M.F., Strobl, D.C., Zappia, L., Dugas, M., Colomé-Tatché, M., Theis, F.J.: Benchmarking atlas-level data integration in single-cell genomics. Nature Methods 19(1), 41–50 (2022) 10.1038/s41592-021-01336-8

[45] Levenshtein, V.I.: Binary codes capable of correcting deletions, insertions, and reversals. Soviet Physics Doklady 10(8), 707–710 (1966). English translation of the original Russian article published in Doklady Akademii Nauk SSSR, 163(4):845–848, 1965

[46] Burkhard, W.A., Keller, R.M.: Some approaches to best-match file searching. Communications of the ACM 16(4), 230–236 (1973) 10.1145/362003.362025

